# Extracellular matrix integrity regulates GABAergic plasticity in the hippocampus

**DOI:** 10.1101/2024.07.30.605739

**Authors:** Jadwiga Jabłońska, Grzegorz Wiera, Jerzy W. Mozrzymas

## Abstract

The brain’s extracellular matrix (ECM) is crucial for neural circuit functionality, synaptic plasticity, and learning. While the role of the ECM in excitatory synapses has been extensively studied, its influence on inhibitory synapses, particularly on GABAergic long-term plasticity, remains poorly understood. This study aims to elucidate the effects of ECM components on inhibitory synaptic transmission and plasticity in the hippocampal CA1 region. We focus on the roles of chondroitin sulfate proteoglycans (CSPGs) and hyaluronic acid in modulating inhibitory postsynaptic currents (IPSCs) at two distinct inhibitory synapses formed by somatostatin (SST)-positive and parvalbumin (PV)-positive interneurons onto pyramidal cells (PCs). Using optogenetic stimulation in brain slices, we observed that acute degradation of ECM constituents by hyaluronidase or chondroitinase-ABC did not affect basal inhibitory synaptic transmission. However, short-term plasticity, particularly burst-induced depression, was enhanced at PV→PC synapses following enzymatic treatments. Long-term plasticity experiments demonstrated that CSPGs are essential for NMDA-induced iLTP at SST→PC synapses, whereas the digestion of hyaluronic acid by hyaluronidase impaired iLTP at PV→PC synapses. This indicates a synapse-specific role of CSPGs and hyaluronic acid in regulating GABAergic plasticity. Additionally, we report the presence of cryptic GABAergic plasticity at PV→PC synapses induced by prolonged NMDA application, which became evident after CSPG digestion and was absent under control conditions. Our results underscore the differential impact of ECM degradation on inhibitory synaptic plasticity, highlighting the synapse-specific interplay between ECM components and specific GABAergic synapses. This offers new perspectives in studies on learning and critical period timing.

## Introduction

The extracellular matrix (ECM) of the brain, a complex network of molecules, is essential for neural circuit functionality, synaptic plasticity, and learning. Although extensive research has addressed the role of ECM in modulating excitatory synaptic transmission and memory formation [1], [2], its influence on inhibitory synapses and their plasticity remains poorly understood. This study aims to explore the role of ECM components in the regulation of inhibitory synaptic transmission and GABAergic long-term plasticity.

All central nervous system cells are embedded in a diffuse extracellular matrix comprising various molecules, such as hyaluronan, chondroitin sulfate proteoglycans (CSPGs), heparan sulfate proteoglycans, and glycoproteins, including thrombospondins, tenascins, and extracellular proteases [3]. The complex network of ECM regulates the diffusion of neurotransmitters, ions, and metabolites, providing structural and biochemical support to neurons and synapses [4]. Additionally, inhibitory GABAergic parvalbumin-positive (PV) interneurons, and some pyramidal cells, are enwrapped by a specialized, condensed ECM rich in hyaluronic acid, known as perineuronal nets (PNNs). These net-like structures are crucial for regulating synaptic stability and excitatory neuroplasticity [5], [6].

The interplay between ECM constituents and inhibitory GABAergic synapses is becoming increasingly evident. For instance, tenascin R interacts with GABA_B_ receptors, modulating presynaptic activity to increase basal excitatory transmission while reducing perisomatic GABAergic inhibition [7]. Additionally, mice concurrently deficient in brevican, neurocan, tenascin-C, and -R exhibit a synaptic excitatory/inhibitory imbalance, with increased decreased number of inhibitory synapses within the PNNs [8]. Acute modifications of the brain ECM also impact inhibitory function. Chondroitinase ABC (Ch-ABC)-mediated degradation of CSPGs results in decreased inhibitory synaptic puncta and diminished gamma rhythmic activity in the prefrontal [9] and the visual cortex of mice [10]. In vitro studies further support the role of the ECM in maintaining inhibitory network stability, demonstrating that ECM digestion reduces inhibitory connectivity [11]. Similar Ch-ABC-induced reductions in inhibitory activity have been observed in the medial entorhinal cortex, which were accompanied by dysfunction in the grid cell network physiology [12]. Additionally, hyaluronan degradation promotes seizure-like activity and enhances learning ability in mice [13], [14], presumably due to reduced synaptic inhibition. The picture emerges that ECM development tends to downregulate inhibition and to temper synaptic plasticity by favoring closure of critical windows [15]. However, it remains unclear whether ECM modifications that downregulate inhibition affect also synapse-specific GABAergic long-term plasticity.

Recent studies have highlighted the crucial interplay between learning, inhibitory synapses, and the ECM. Learning modifies the levels of perisomatic inhibition in engram neurons [16], [17], and this process relies on the extracellular matrix [17]. PNNs are crucial for brain maturation and synapse stabilization, particularly in safeguarding long-term memory by restricting excessive plasticity [18]. This protective mechanism hinges on synaptic inhibition, as evidenced by Ch-ABC-treated brain slices showing inhibitory long-term potentiation (iLTP) in response to theta-burst stimulation, unlike the inhibitory long-term depression (iLTD) observed in controls [18]. Furthermore, enzymatic degradation of CA1 PNNs with Ch-ABC can revert memory to a juvenile-like state, characterized by immature parvalbumin-positive (PV) inhibitory circuitry and imprecise memory formation [17]. Therefore, understanding how the extracellular milieu interacts with diverse inhibitory synapses is vital for gaining comprehensive insights into the role of brain ECM in learning and memory formation.

In this study, we investigated the cellular mechanisms underlying the roles of chondroitin sulfate proteoglycans and hyaluronic acid in inhibitory synaptic transmission and plasticity within the hippocampal CA1 region. To this end, we recorded optogenetically evoked IPSC originating from two distinct GABAergic inputs to CA1 pyramidal cells, namely from somatostatin (SST)-positive and parvalbumin (PV)-positive interneurons. Our findings reveal that these ECM components play synapse-specific roles during NMDA-induced GABAergic plasticity. Specifically, our results indicate that the degradation of ECM constituents can constrain plasticity at one type of inhibitory synapse, while enhancing it at another. Altogether, our results indicate that ECM degradation leads to distinct modifications in synaptic plasticity (both short-term and long-term), depending on the type of interneuron innervating the pyramidal cell.

## Materials and Methods

### Ethical approval, animals and reagents

All procedures involving the use of animals were conducted in accordance with the Act on the Protection of Animals Used for Scientific or Educational Purposes in Poland (Act of January 15, 2015, with subsequent amendments) and EU Directive 2010/63/EU. The Polish Ministry of the Environment has granted approval for experiments involving genetically modified organisms (decision numbers 144/2018 and 69/2023). The mouse strains used in this study: Sst-IRES-Cre (Jackson Labs stock #013044), Pvalb-IRES-Cre (Jackson Labs stock #017320), Ai32 (B6.Cg-Gt(ROSA)26Sortm32(CAG-COP4*H134R/EYFP)Hze/J; Jackson Labs stock #024109), C57BL/6 parental lines, and their offspring (SST-Cre::Ai32 and PV-Cre::Ai32). These mice were maintained under a 12-hour dark/light cycle and provided food ad libitum.

Electrophysiological experiments were performed on acute brain slices from male and female offspring of SST-Cre::Ai32 and PV-Cre::Ai32 crossings with interneuron-specific expression of channelrhodopsin-2. Immunofluorescence staining was performed on tissues from C57BL/6 mice. The key reagents used are listed in Table 1. Reagents that were not listed were purchased from Merck (Sigma-Aldrich).

**Table 1.** Suppliers and catalog numbers of key reagents.

| Reagent | Source | ID |
| --- | --- | --- |
| Anti-integrin β1 (monoclonal) | Merck | Catalog #MAB2252 |
| Anti-integrin β3 (monoclonal) | Merck | Catalog #MAB1957 |
| Anti-integrin β1, active conformation (monoclonal) | Merck | Catalog #MAB2079Z |
| Anti-LIBS2 epitope (integrin β3 subunit), active conformation (monoclonal) | Merck | Catalog #MABT27 |
| Anti-vGAT cytoplasmic domain (polyclonal) | Synaptic Systems | catalog #131002 |
| Goat anti-mouse IgG, Alexa Fluor 488 | Thermo Fisher Scientific | Catalog #A-11029 |
| Goat anti-rabbit IgG, Alexa Fluor 633 | Thermo Fisher Scientific | Catalog #A-21071 |
| WFA, biotin conjugate | Sigma-aldrich | Catalog #L1516 |
| Streptavidin, alexa488 conjugate | Thermo Fisher Scientific | Catalog #S11223 |
| Gelatin | VWR Chemicals | Catalog #24350-262 |
| Normal horse serum | Vector Laboratories | Catalog #S-2000 |
| Fluoroshield with DAPI | Sigma-Aldrich | Catalog #F6057 |
| hyaluronidase | Sigma-Aldrich | Catalog #H1136 |
| Chondroitinase ABC | Sigma-Aldrich | Catalog #C3667 |

### Slices preparation

Acute hippocampal slices were prepared from P60-90 mice of all genotypes, according to the protocol described by Ting et al. [19], with minor modifications. Brains were removed from mice anesthetized with 5% isoflurane and placed in ice-cold holding aCSF (in mM: 92 NaCl, 2.5 KCl, 1.2 NaH_2_PO_4_, 30 NaHCO_3_, 20 HEPES, 25 D-glucose, 5 sodium ascorbate, 2 thiourea, 3 sodium pyruvate, 1.3 MgSO_4_, 2.5 CaCl_2_), bubbled with 95% O_2_ and 5% CO_2_. The brains were then cut into 350 μm transverse slices in the same solution using a vibratome (VT1200S, Leica). The slices were then transferred to recovery aCSF (in mM: 93 N-methyl-D-glucamine, 2.5 KCl, 1.2 NaH_2_PO_4_, 30 NaHCO_3_, 20 HEPES, 25 glucose, 5 sodium ascorbate, 2 thiourea, 3 sodium pyruvate, 10 MgSO_4_, 0.5 CaCl_2_) heated to 34°C. Increasing volumes from 0.25 to 2ml of 2M NaCl ware added over 25-30 min. incubation period. Procedure allows avoidance of rapid NaCl concentration changes, that may damage cell membranes and thus interfere with seal formation in patch-clamp (Ting et al. 2018). After recovery, the slices were transferred to room temperature holding aCSF and used for experiments within 8 hours.

### Enzymatic treatment

To degrade hyaluronic acid (HA) or chondroitin sulfate proteoglycans (CSPG) before analyzing GABAergic synaptic transmission and plasticity, individual slices were incubated for 2 hours at room temperature in 2.3 ml of holding aCSF supplemented with hyaluronidase from *Streptomyces hyalurolyticus* (Sigma #H1136; 70 U/ml) or protease-free chondroitinase ABC from *Proteus vulgaris* (Sigma #C3667, 0.25 U/ml), and bubbled with carbogen. Hyaluronidase cleaves the β-GlcNAc-[1-4] glycosidic bonds in HA, producing tetra- and hexasaccharides, whereas chondroitinase cleaves 1,4-hexosaminidic bonds in chondroitin sulfate and dermatan sulfate, resulting in di- and tetrasaccharides. Wisteria floribunda agglutinin (WFA) was used to visualize perineuronal nets and further confirm enzymatic digestion. Immediately after treatment, the slices were placed in a submerged recording chamber with continuous superfusion of aCSF that did not contain the enzyme. Control slices were incubated under identical conditions in 2.3 ml of holding aCSF without enzyme, serving as sham controls.

### Electrophysiological recordings

The hippocampal slices were placed in a recording chamber perfused at 2.5-3 ml/min with carbogen-saturated aCSF (in mM: 119 NaCl, 26.3 NaHCO_3_, 11 glucose, 2.5 KCl, 1 NaH_2_PO_4_, 1.3 MgSO_4_, 2.5 CaCl_2_, pH 7.4). Borosilicate glass electrodes with a resistance of 3-5 MΩ, were filled with an intracellular solution containing 135 mM K-gluconate, 5 mM NaCl, 10 mM HEPES, 5 mM MgATP, 0.3 mM GTP, and 10 mM phosphocreatine. Whole-cell patch clamp measurements were recorded at 28°C using a MultiClamp 700B amplifier and Digidata 1550B digitizer (Molecular Devices). The recorded signals were filtered at 6 kHz and digitalized at 20 kHz. To preserve physiological chloride balance, the internal pipette solution contained a low concentration of Cl^-^ ions. For each cell, the reversal potential of synaptic GABA_A_ receptors was measured by recording the IPSCs at different holding potentials from -90 to -55 mV. The hyperpolarizing IPSCs were recorded at -60mV holding potential. The liquid junction potential was not corrected, and series resistance was not compensated during the recordings.

Pyramidal cells (PCs) were identified morphologically based on their soma placement within the stratum pyramidale and electrophysiologically based on their firing patterns. Only neurons with regularly accommodating action potentials and noticeable sags were considered PCs. Input resistance was monitored throughout the recordings and experiments showing >20% divergence were excluded from the analysis.

GABAergic long-term synaptic plasticity was induced by briefly exposing slices to 20 μM NMDA in bath solution for 1:15-2:30 minutes. No additional substances were added during the experiments. Each recording was conducted for at least 10 minutes before and 30 minutes after NMDA administration. The magnitude of long-term GABAergic plasticity was calculated as percent change, comparing the average amplitude of IPSCs between the baseline period (6 minutes before NMDA) and 25-30 minutes post-NMDA administration.

To investigate the short-term plasticity of IPSCs, we used paired-pulse stimulation, delivering pairs of stimuli at various interstimulus intervals (50 ms, 200 ms, 500 ms, 2 s, and 10 s). The paired-pulse ratio was determined by dividing the amplitude of the second IPSC by that of the first. Additionally, to evaluate the burst response, trains of eight stimuli were applied at 200 ms intervals. The burst depression was quantified by calculating the ratio of the average amplitude of the 6th to 8th responses to the amplitude of the first response within each burst.

### Optogenetic stimulation

To activate the cells and axons expressing channelrhodopsin-2, a fiber-coupled DG-4 (Sutter Instruments) equipped with an HQ470/40x excitation filter (Chroma) was utilized. Blue light was directed through the back aperture of a 40x, 1.0 NA microscope objective (Zeiss) at a power density of 5–10 mW/mm^2^. Inhibitory synaptic transmission was evoked using 2 ms light pulses. To maximize the specificity of light excitation, two distinct methods were employed to stimulate the PV- and SST-positive presynaptic interneurons. To evoke SST→PC synaptic transmission, the objective was centered over the stratum oriens, with the aperture fully open. Conversely, for the PV→PC inhibitory transmission, the objective aperture was reduced to limit the light-spot diameter over the stratum pyramidale. This approach allowed for precise and spatially specific stimulation of two different inhibitory inputs to CA1 pyramidal cells.

### Immunofluorescence stainings

Enzyme-treated and control tissues were fixed overnight at 4°C in a solution containing 4% paraformaldehyde, 0.2% picric acid, and phosphate-buffered saline (PBS). The following day, the tissues were rinsed with PBS. Fixed hippocampal slices were then embedded in 15-17% gelatin and sectioned into 40 μm slices using a vibratome (VT1000S, Leica). Free-floating sections were permeabilized with 0.1% Triton-X in PBS and subsequently blocked with 5% normal horse serum (NHS) in PBS-Triton-X for 1 h at room temperature. The impact of ECM digestion on total integrin β1 and β3 levels and their activation was determined using antibodies, as described in a recent study [20]. Primary antibodies (1:100 dilution for anti-total integrin β1, β3, and activated integrin β3; activated integrin β3 1:200; vGAT: 1:500) or Wisteria floribunda agglutinin (WFA; 1:1000) were diluted in 3% NHS in PBS and incubated with the sections for 24 h at 4°C with gentle stirring. Next, sections were incubated with appropriate secondary antibodies and streptavidin (all at 1:1000) for another 24 h at 4°C with gentle stirring. Finally, the sections were mounted on microscope slides using Fluoroshield with DAPI.

### Image acquisition and analysis

The WFA-stained slices were visualized using an Olympus Fluoview 1000S confocal microscope at the Wroclaw University, employing a 20x objective. Z-stacks consisting of three images with a 1 μm step size were scanned in sequential mode. Subsequent image analysis was performed using the Fiji software [21] on maximum intensity projections. The fluorescence integrated density was quantified separately for the stratum radiatum and combined stratum pyramidale and stratum oriens.

Sections immunostained for integrins were imaged using a 60x oil immersion objective. Z-stacks comprising six images at 0.95 μm intervals were acquired in sequential mode. Further image analysis was conducted using Fiji software to create maximum-intensity projections. The vicinity of the inhibitory synapses was specified by creating a mask (ROI) of vGAT-positive puncta, extended by two pixels in every direction. The hippocampal layers were manually outlined. The mean fluorescence values of the integrin signal in the vGAT-positive areas were measured and normalized to those of the control group within each batch of staining. Thresholded images of integrin staining and vGAT-positive ROIs were used in the particle analysis to determine the size and count of integrin puncta. Eight independently treated slices for each condition (Hyase, Ch-ABC, and sham-treated control) from four mice were used for the statistical analysis.

### Quantification and statistical analysis

Several predefined criteria were followed in the data analysis process: (1) no outliers were excluded from the dataset, (2) random assignment of slices to treatments was implemented, (3) investigators were not blinded to the treatment during recording and analysis, and (4) minimal sample sizes were determined through power analysis, aiming for a minimum power level of 0.80.

Electrophysiological recordings were analyzed using the Clampfit 10.7 software. The onset kinetics of the IPSC were assessed based on the 10% to 90% rise time, while the decay phase was analyzed by fitting a biexponential function:*y*(*t*) = *A*_1_*exp*(−*t*/*τ*_*fast*_) + *A*_2_*exp*(−*t*/*τ*_*slow*_), where τ_fast_ and τ_slow_ represent the time constants, and A_1_ and A_2_ denote the amplitudes of the fast and slow components, respectively. The mean decay time constant (τ_mean_) was calculated using the formula *τ*_*mean*_ = *a*_1_*τ*_*fast*_ + *a*_2_*τ*_*slow*_, where *a*_1_ = *A*_1_/(*A*_1_ + *A*_2_) and *a*_2_ = *A*_2_/(*A*_1_ + *A*_2_).

All statistical analyses were conducted using the SigmaPlot 11 software. Each experimental condition included independent experiments using slices from at least 4 different mice. Normality of the datasets was assessed using the Shapiro-Wilk test, and comparisons between groups were performed using two-tailed paired or unpaired Student *t*-tests or Mann-Whitney U tests, as appropriate. The size of the analyzed groups and the specific statistical tests performed are indicated in the figure legends. Statistical significance was denoted as follows: NS (non-significant) p > 0.05; * p < 0.05; ** p < 0.01; and *** p < 0.001.

## Results

### Enzymatic digestion of brain extracellular matrix

To verify the efficiency of ECM digestion, we labelled perineuronal nets (PNNs) with Wisteria floribunda agglutinin (WFA) in chondroitinase-, hyaluronidase-, and sham-treated hippocampal slices (Fig. 1A). WFA staining revealed a distribution pattern of chondroitin sulfate proteoglycans (CSPGs) typical of the adult hippocampal CA1 region, with moderate staining of the neuropil in the stratum radiatum and stratum lacunosum moleculare and intense staining of PNNs in the stratum oriens and stratum pyramidale [22].

**Figure 1.**
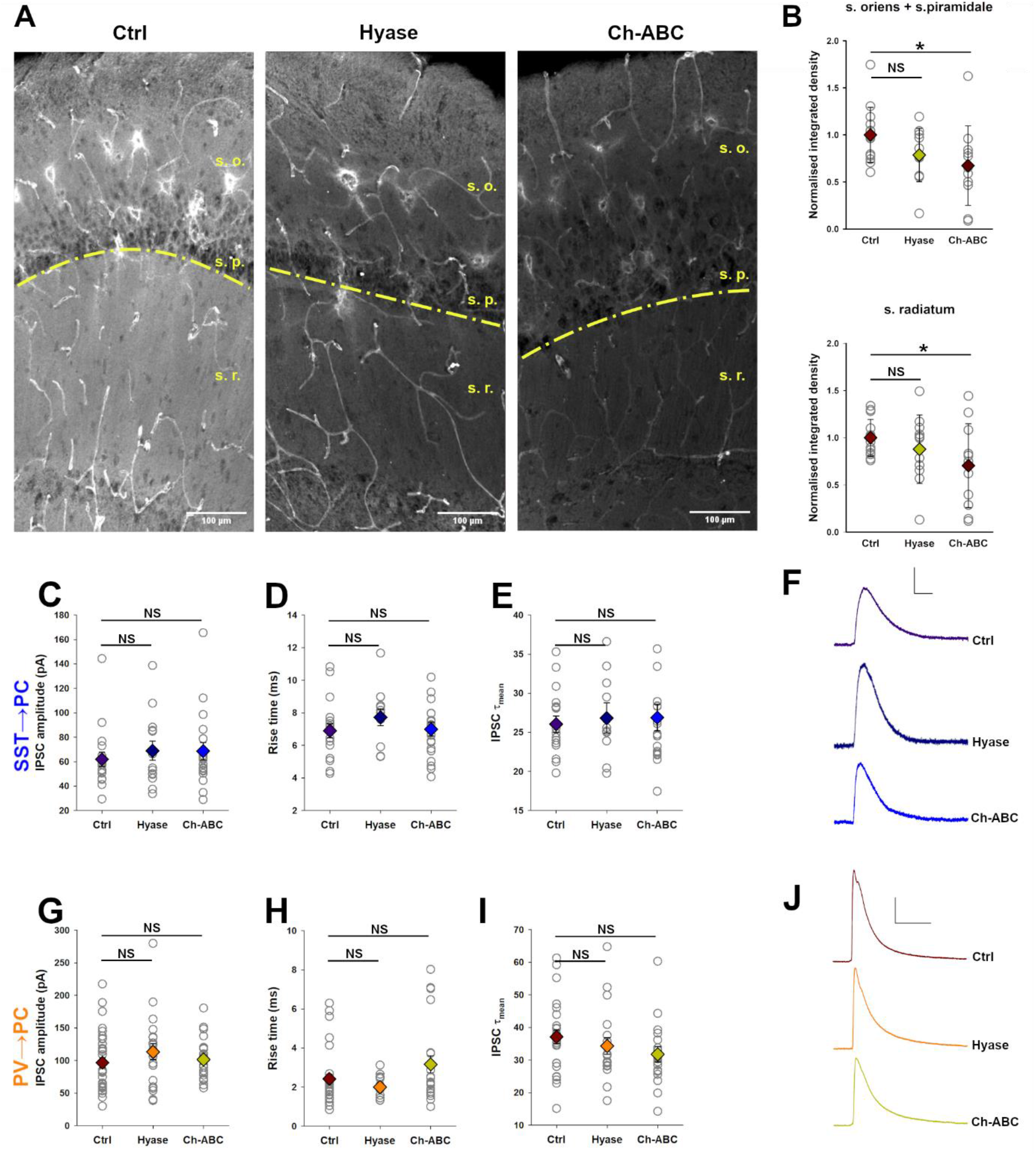
Effect of hyaluronan and chondroitin sulfate digestion on WFA staining intensity and optogenetically evoked IPSC properties. **A**. Representative images of untreated, hyaluronidase-treated, and chondroitinase ABC-treated slices stained with Wisteria floribunda agglutinin (WFA). **B**. WFA fluorescence integrated density in specific hippocampal regions: combined stratum oriens and stratum pyramidale (top) and stratum radiatum (bottom); (Ctrl n=14, Hyase n=11, Ch-ABC n=11). **C-E**. Analysis of IPSCs recorded in inhibitory synapses between somatostatin (SST) interneurons and CA1 pyramidal cells in sham-treated slices (Ctrl) or slices treated with hyaluronidase (Hyase) or chondroitinase (Ch-ABC). (**C**) Comparison of IPSC amplitudes (Ctrl n=18, Hyase n=14, Ch-ABC n=19), (**D**) Comparison of rise times (Ctrl n=17, Hyase n=12, Ch-ABC n=19). (**E**) Comparison of decay time constants (τ_mean_) (Ctrl n=17, Hyase n=11, Ch-ABC n=18). **F**. Representative IPSCs recorded at SST→CA1 PC synapses in control (top), hyaluronidase-treated (middle), and chondroitinase-treated (bottom) slices (scale: 20 pA, 200 ms). **G-I**. Analysis of IPSCs recorded in inhibitory synapses between parvalbumin (PV) interneurons and CA1 pyramidal cells in sham-treated slices (Ctrl) or slices treated with hyaluronidase or chondroitinase (Ch-ABC). (**G**)Comparison of IPSC amplitudes (Ctrl n=34, Hyal n=21, Ch-ABC n=22) (**H**) Comparison of rise times (Ctrl n=33, Hyase n=18, Ch-ABC n=22). (**I**) Comparison of decay time constants (Ctrl n=27, Hyase n=19, Ch-ABC n=18). **J**. Representative IPSCs recorded at the PV→CA1 PC synapses in control (top), hyaluronidase-treated (middle), and chondroitinase-treated (bottom) slices (scale: 40 pA, 40 ms). Data are presented as mean ± SEM. Statistical significance was determined by the t-test vs. control group: *p < 0.05, ns: not significant.

Slices treated with chondroitinase showed reduced PNN staining intensity across all analyzed layers within the CA1 region (stratum radiatum: control normalized fluorescence intensity: 1.0±0.19, Ch-ABC: 0.7±0.44, p=0.046; stratum oriens: control: 1.0±0.29, Ch-ABC: 0.67±0.42, p=0.032) (Fig. 1B). The reduced fluorescence intensity confirmed the efficacy of the digestion conditions [12], [23], [24]. Although hyaluronidase treatment resulted in a slight but noticeable decrease in the WFA signal, these changes were not statistically significant (stratum radiatum Hyase: 0.87±0.36, p=0.29 vs. control; stratum oriens Hyase: 0.79±0.28, p=0.079; Fig. 1B). The lower efficiency of hyaluronidase on PNNs and CSPGs compared to chondroitinase ABC treatment can be attributed to differences in substrate specificity, which is consistent with previous findings [5].

### Enzymatic digestion of ECM does not alter basal inhibitory synaptic transmission in the hippocampus

To investigate the impact of ECM removal on inhibitory synaptic transmission, we optogenetically stimulated two distinct GABAergic inputs into hippocampal CA1 pyramidal cells (PCs). As expected, owing to electrotonic filtering, optogenetically evoked IPSCs at PV→PC synapses, which are primarily located on the cell body and proximal dendrites of CA1 PCs, exhibited faster kinetics than those at SST→PC synapses, which are primarily located on the dendritic tree (Figs. 1C-J) [25]. We then compared the amplitude and kinetics of both inhibitory inputs between slices treated with hyaluronidase, chondroitinase, or sham-treated control slices (Figs. 1C-J). We did not observe any significant changes in IPSC amplitude in either SST→PC synapses (Ctrl: 61.7±5.87 pA, Hyase: 68.9±7.84 pA p=0.47, Ch-ABC: 68.6±7.15 pA p=0.48; Figs. 1C,F) or PV→PC synapses (Ctrl: 96.8±7.66 pA; Hyase: 113.4±12.1 pA, *t* test vs. Ctrl p=0.23; Ch-ABC: 101.5±6.89 pA p=0.67; Figs. 1G,J). Similarly, enzymatic treatment did not affect the kinetics of evoked IPSC in GABAergic projections. Both the rise time (SST→PC Ctrl: 6.9±0.4 ms, Hyase: 7.7±0.5 ms p=0.99, Ch-ABC: 7.0±0.4 ms p=0.31; PV→PC Ctrl: 2.4±0.2, Hyase: 2.0±0.1 ms p=0.21, Ch-ABC: 3.2±0.4 ms p=0.1; Figs, 1D,H) and decay τ_mean_ kinetics (SST→PC Ctrl: 26.0±1.1 ms, Hyase: 26.8±1.9 ms p=0.85, Ch-ABC: 26.9±1.7 ms p=0.79; PV→PC Ctrl: 37.1±2.1 ms, Hyase: 34.3±2.6 ms p=0.4, Ch-ABC: 31.8±2.4 ms p=0.1; Figs. 1E,I) remained unaffected. Therefore, acute degradation of brain hyaluronic acid or CSPGs does not alter the basal properties of inhibitory synaptic transmission between somatostatin or parvalbumin interneurons and CA1 pyramidal cells.

In excitatory synapses, enzymatic digestion of ECM-derived perisynaptic compartments, which restricts the surface diffusion of desensitized AMPA receptors [26], accelerates receptor diffusion and suppresses short-term synaptic plasticity [27]. Analogous mechanisms have not been systematically studied at inhibitory synapses, although it is known that the lateral movement of synaptic inhibitory receptors can be influenced by ECM and adhesion receptors [28] or the extracellular protease MMP3 [29]. Therefore, we investigated the impact of ECM digestion on short-term synaptic plasticity at different inhibitory synapses. In our experiments, we used two distinct protocols to assess whether the degradation of HA or CSPGs affected the inhibitory short-term plasticity of light-evoked inhibitory postsynaptic currents. First, paired pulses were applied at intervals of 50 ms, 200 ms, 500 ms, 2 s, and 10 s. Neither the SST→CA1PC nor PV→CA1 PC synapses exhibited significant changes following enzyme treatment (Fig. S1). Second, burst stimulation with eight pulses applied at 5 Hz was used to test the prolonged synapse activation. In all recordings, we observed burst-induced depression of IPSC, a form of short-term plasticity. We analyzed this effect by calculating the ratio between the mean of the IPSC amplitudes of the last stimuli within the burst and the first one. The burst-induced short-term depression in SST→PC synapses did not change after ECM digestion (Ctrl: 0.61±0.02, Hyase: 0.65±0.04, p=0.22; Ch-ABC: 0.62±0.02, p=0.65) (Figs. 2A-C). In contrast, PV→PC transmission showed enhanced burst-induced depression after both enzymatic treatments, with a larger effect observed upon chondroitinase digestion (Ctrl: 0.42±0.02, Hyase: 0.37±0.02, p=0.05; Ch-ABC: 0.33±0.02, p=0.0007 vs. Ctrl; Figs. 2D-F). This finding aligns with the prevalence of perineuronal nets (PNNs) around PV-positive interneurons, and suggests a higher sensitivity of these neurons to ECM manipulation.

**Figure 2.**
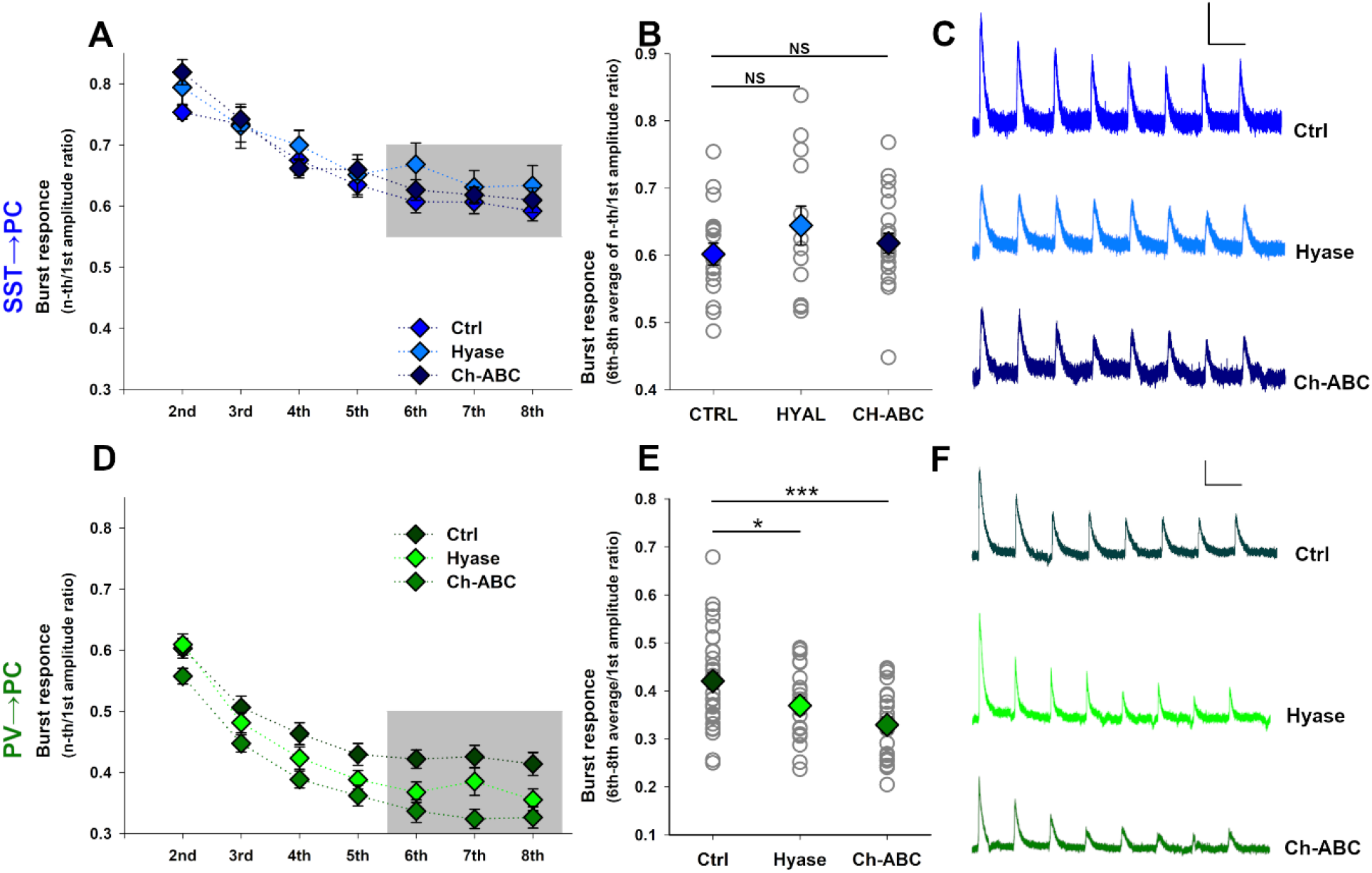
Effects of enzymatic digestion of hyaluronic acid and chondroitin sulfate proteoglycans on short-term plasticity of inhibitory synaptic transmission in the CA1 field of the hippocampus. **A**. Burst-induced depression at the SST→CA1 pyramidal cell (PC) synapses. Average ratios of the amplitude of the n^th^ response to the first inhibitory postsynaptic current (IPSC) within a burst are shown for sham-treated slices (Ctrl) and slices treated with chondroitinase ABC (Ch-ABC) or hyaluronidase (Hyase). The gray area indicates the data used to calculate the burst-induced depression parameter. **B**. Summary plot of burst-induced depression recorded in the analyzed groups. This parameter was calculated as the ratio of the mean averaged IPSC amplitude of the 6^th^-8^th^ responses within a burst to the amplitude of the 1^st^ IPSC (Ctrl n=18, Hyase n=13, Ch-ABC n=22). **C**. Representative traces showing burst-induced depression at the SST→CA1 PC synapses in control (top), hyaluronidase-treated (middle), and chondroitinase-treated (bottom) slices (scale: 20 pA, 200 ms). **D**. Burst-induced depression at PV→CA1 PC synapses in sham-treated slices (Ctrl) and slices treated with Chondroitinase ABC (Ch-ABC) or hyaluronidase (Hyase). **E**. Summary plot of burst-induced depression recorded in the analyzed groups (Ctrl n=35, Hyase n=21, Ch-ABC n=21). **F**. Representative traces showing burst-induced depression at PV→CA1 PC synapses in control (top), hyaluronidase-treated (middle), and chondroitinase-treated (bottom) slices (scale: 20 pA, 200 ms). All data are presented as mean ± SEM. Statistical significance was determined by the t-test vs. the control group: *p < 0.05, ***p < 0.001, ns – not significant.

### NMDA-induced iLTP in SST→CA1 PC synapses requires chondroitin sulfate proteoglycans but not hyaluronic acid

Given the active involvement of extracellular matrix constituents in the molecular mechanisms underlying long-term plasticity at excitatory synapses [5], [24], [30], we sought to investigate the roles of hyaluronic acid (HA) and chondroitin sulfate proteoglycans (CSPGs) in long-term GABAergic plasticity. Specifically, we focused on inhibitory plasticity induced in SST→ CA1 PC projection by a brief application of 20 μM NMDA for 1 min and 45 s.

Recordings from control sham-treated slices revealed a stable increase in IPSC amplitude 24-26 minutes after NMDA application (53.6±3.2 pA before, 66.1±4.8 pA after, p=0.002, paired *t*-test; Figs. 3A-D). In slices treated with hyaluronidase (Hyase), NMDA-iLTP was expressed at similar level to that in control conditions (71.8±10.5 pA before, 87.3±14.4 pA after, p=0.02, paired *t*-test; Figs. 3A,C,D). This suggests that the acute degradation of hyaluronan does not significantly affect iLTP in SST→PC synapses (Ctrl: 125.9±7.6% of baseline IPSC amplitude, Hyase: 120.0±5.6%, p=0.56; *t*-test; Fig. 3B). Conversely, chondroitinase ABC treatment markedly impaired the induction of NMDA-iLTP in SST→PC synapses (71.5±8.3 pA before, 77.5±10.8 pA after, p=0.12; Figs. 3A-C), significantly reducing the magnitude of IPSC potentiation (Ch-ABC: 106.1±4.5%, p=0.03; *t*-test vs. Ctrl; Fig. 3B). These observations confirm that CSPGs, but not hyaluronic acid, are crucial for the induction of long-term plastic changes at synapses between SST interneurons and the dendrites of CA1 pyramidal cells, underscoring the specific involvement of CSPGs in the regulation of GABAergic plasticity.

**Figure 3.**
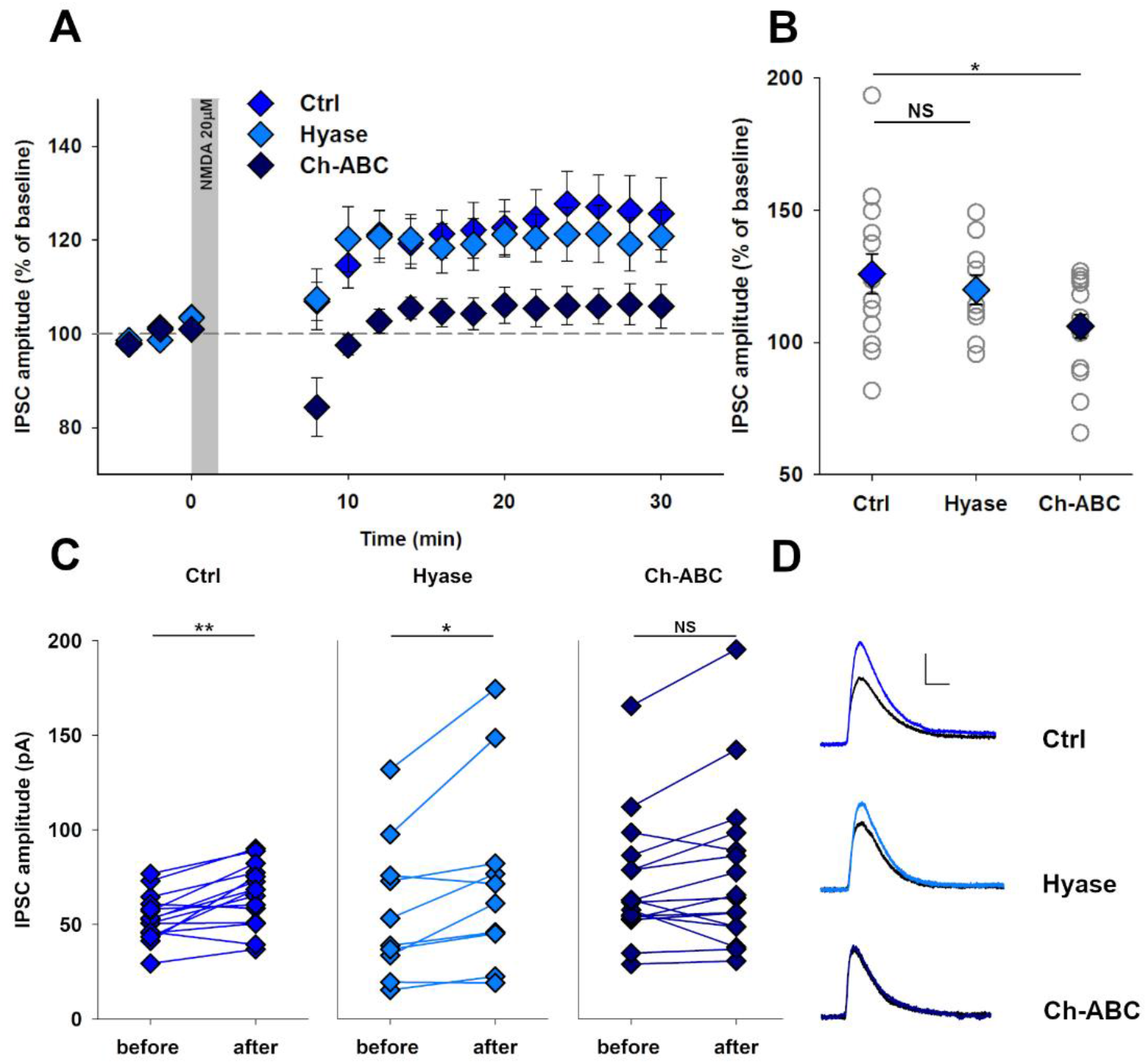
Chondroitin sulfate, but not hyaluronic acid, digestion impairs NMDA-induced iLTP in inhibitory synapses between CA1 somatostatin (SST) interneurons and pyramidal cells (PC) **A**. GABAergic long-term potentiation (iLTP) induced at SST→PC synapses by bath application of NMDA for 1 min 45 s, recorded in sham-treated slices, and slices treated for 2 hours with hyaluronidase (Hyase) or chondroitinase ABC (Ch-ABC). The gray area indicates the duration of NMDA application. **B**. Summary plot of iLTP magnitudes assessed 24-30 minutes after induction in SST→PC synapses (Ctrl n=15, Hyase n=10, Ch-ABC n=15). Diamonds represent mean ± SEM. Statistical significance was determined by the t-test vs. control group. **C**. Comparison of IPSC amplitudes in SST→PC synapses recorded before and 26-30 minutes after iLTP induction with NMDA. Statistical significance was determined using the paired t-test. **D**. Representative traces showing IPSCs at the PV→CA1 PC synapses recorded before (black) and 25-30 minutes after iLTP induction (blue) in control (top), hyaluronidase-treated (middle), and chondroitinase-treated (bottom) slices (scale: 20 pA, 20 ms). *p < 0.05, **p < 0.01, ns – not significant.

### ECM digestion affects integrins

A previous study showed that CSPG digestion promotes the activation of integrins, receptors for ECM constituents, and enhances the structural dynamics of dendritic spines [22]. Thus, we asked whether ECM digestion could affect the level of integrin activation. As brain integrins can be broadly divided into containing β1 or β3 subunit, we assessed whether ECM digestion affects the level of total pool of these integrins (Figs. 4A,C), and the level of their active forms adopting an extended conformation (Figs. 4B,D).

**Figure 4.**
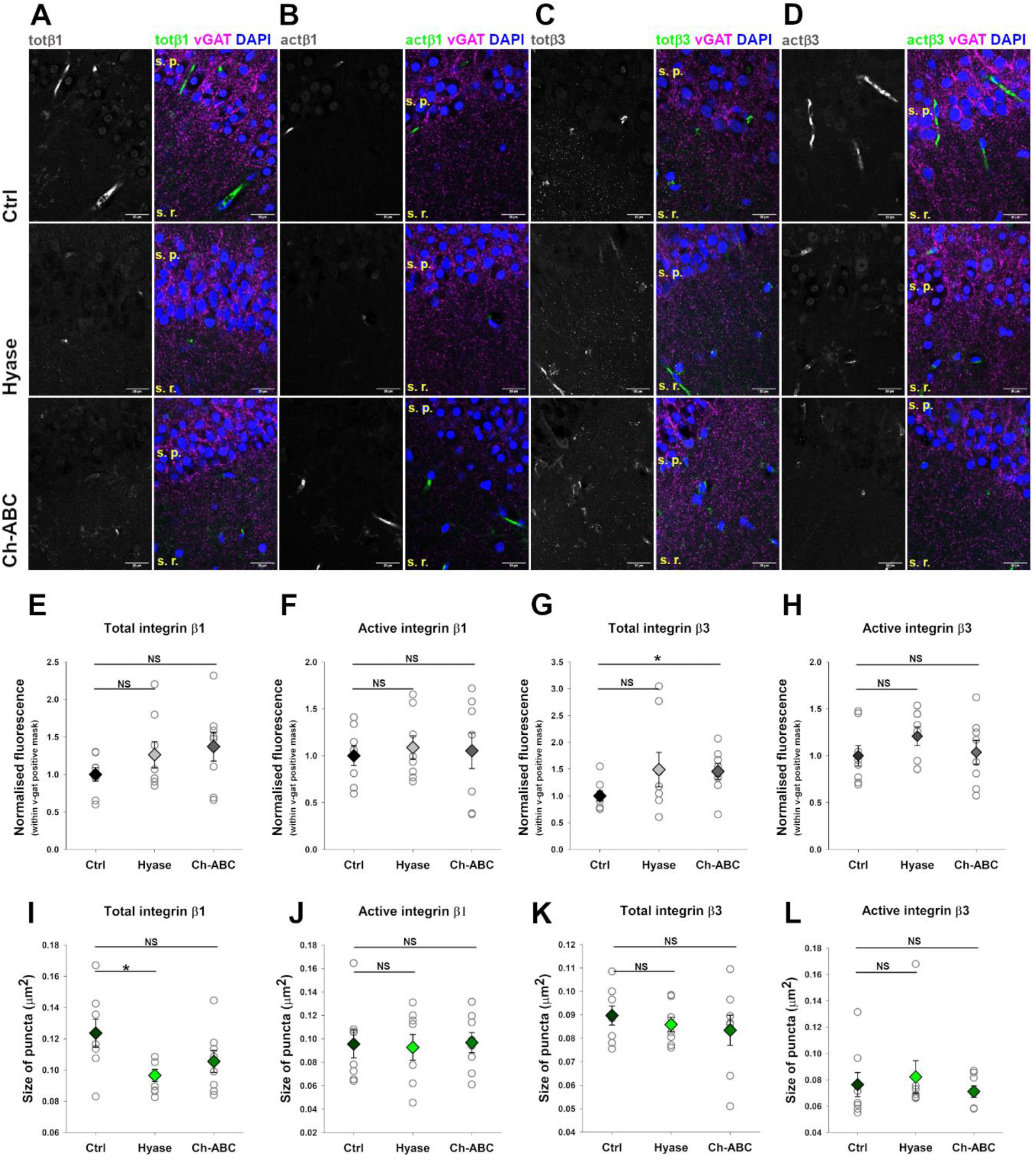
Impact of hyaluronan and chondroitin sulphate degradation on the levels of total pool and active forms of integrin β1 and β3. **A-D**. Representative images of sham-treated (Ctrl), hyaluronidase-treated (Hyal), and chondroitinase ABC-treated (Ch-ABC) slices stained with DAPI (cyan), vGAT (red), and different pools of integrins (green). The slices were stained with antibodies targeting the total pool of integrin β1 (**A**), extended conformation of integrin β1 (presumably active) (**B**), total pool of integrin β3 (**C**), and extended conformation of integrin β3 (presumably active) (**D**). **E-H**. Analysis of the average fluorescence intensity of integrin puncta near inhibitory synapses identified by vGAT staining, in the CA1 stratum radiatum. **I-K**. Analysis of the average size of integrin puncta near inhibitory synapses, visualizes using vGAT staining, in the CA1 stratum radiatum. Data represent n=8 independently treated slices from 4 mice. Colored diamonds indicate mean ± SEM. Statistical significance was determined by t-test vs. Ctrl group: *p < 0.05, ns – not significant. Scale bars 20μm.

Treatment with either hyaluronidase or chondroitinase showed a tendency to increase the fluorescence intensity of the total pool of integrin β1 in CA1 stratum radiatum, although these changes were not statistically significant (Ctrl: 1.0 ± 0,0924, Hyase: 1,26± 0,18, p = 0.21; Ch-ABC: 1,37± 0,19, p = 0.1 vs. Ctrl; Fig. 4E). Similarly, staining for the active extended conformation of integrin β1 did not reveal any significant differences (Ctrl: 1.0 ± 0,11, Hyse: 1.09 ± 0,13, p = 0.61; Ch-ABC: 1.05 ± 0,19, p = 0.81 vs. Ctrl; Fig. 4F). For the total pool of integrin β3, both treatments showed a tendency to increase fluorescence intensity, with a statistically significant increase observed only after chondroitinase treatment (Ctrl: 1.0 ± 0.1, Hyase: 1.49 ± 0.32, p = 0.28; Ch-ABC: 1.46 ± 0.15, p = 0.02 vs. Ctrl; Fig. 4G). However, no significant changes were detected in the level of the active form of integrin β3 (Ctrl: 1.0 ± 0.11, Hyase: 1.21 ± 0.1, p = 0.18; Ch-ABC: 1.04 ± 0.13, p = 0.66 vs. Ctrl; Fig. 4H).

Despite the lack of significant changes in fluorescence intensity of total pool of integrin β1, additional analysis revealed an increased number and decreased size of integrin puncta after hyaluronidase treatment (puncta density per µm^2^: Ctrl: 8.49 ± 0.65, Hyase: 10.54 ± 0.44, p=0.02; Ch-ABC: 10.01 ± 0.64, p=0.12; Fig. 2A; area of puncta: Ctrl: 0.12 ± 0.01 µm^2^, Hyase: 0.097 ± 0.004 µm^2^, p=0.01; Ch-ABC: 0.105 ± 0.007 µm^2^, p=0.13; Fig. 4I), suggesting dispersion of total integrin β1 clusters due to ECM degradation. No significant differences ware noted for the active form of integrin β1 (puncta density per µm^2^: Ctrl: 11.8 ± 1.2, Hyase: 12.3 ± 1.7, p=0.83; Ch-ABC: 11.1 ± 1.1, p=0.66; Fig. S2B; area of puncta: Ctrl: 0.095 ± 0.012 µm^2^, Hyase: 0.093 ± 0.011 µm^2^, p=0.86; Ch-ABC: 0.097 ± 0.0086 µm^2^, p=0.93; Fig. 4J), total pool of integrin β3 (puncta density per µm^2^: Ctrl: 11.5 ± 0.52, Hyase: 11.8 ± 0.39, p=0.65; Ch-ABC: 12.7 ± 1.2, p=0.34; Fig. S2C; area of puncta: Ctrl: 0.090 ±0.0041 µm^2^, Hyase: 0.086 ± 0.0031 µm^2^, p=0.46; Ch-ABC: 0.083 ± 0.0065 µm^2^, p=0.42; Fig. 4K) nor the active form of integrin β3 (puncta density per µm^2^: Ctrl:14.4 ± 1.18, Hyase: 14.1 ± 0.81, p=0.84; Ch-ABC: 14.4 ± 0.88, p=0.99; Fig. S2D; area of puncta: Ctrl: 0.076 ±0.0092 µm^2^, Hyase: 0.082 ±0.012 µm^2^, p=0.71; Ch-ABC: 0.071 ±0.0044 µm^2^, p=0.61; Fig. 4L). In summary, degradation of extracellular matrix components appears to increase the total levels of β3 integrin in the CA1 stratum radiatum without significantly altering the active forms of both integrin β1 and β3. Thus, changes in integrin activation do not explain the impairments in iLTP observed in the SST→PC synapses after chondroitinase treatment.

### Long-term plasticity in PV→CA1 PC synapses requires hyaluronic acid but not chondroitin sulfate proteoglycans

Our findings indicate that iLTP in dendrite-targeting (SST→PC) inhibitory synapses requires chondroitin sulfate proteoglycans. To further explore the role of ECM in GABAergic plasticity, we analyzed NMDA-induced iLTP in inhibitory synapses located on the soma and proximal dendrites of CA1 pyramidal cells formed by parvalbumin (PV) interneurons. For this purpose, we applied NMDA for 1 min and 15 s. In sham-treated control slices, NMDA application successfully induced iLTP in PV→PC synapses, as evidenced by a significant increase in IPSC amplitude (85.9±8.9 pA before, 97.9±9.3 after, p=0.003, paired *t*-test; Figs. 5A,B). When CSPGs were digested using chondroitinase ABC (Ch-ABC), NMDA application still potentiated PV→PC synapses, with a significant increase in IPSC amplitude (91.8±6.2pA before, 104.1±6.4pA after, p=0.02, paired t-test). However, hyaluronidase (Hyase) treatment abolished NMDA-induced iLTP, as indicated by the lack of a significant change in IPSC amplitude after NMDA treatment (125.4±18.4pA before, 130.5±2.5pA after, p=0.4; Figs. 5C-D). The magnitude of plasticity in comparison to the control group was not different in Ch-ABC-treated slices, but was significantly reduced in the Hyase-treated group (Ctrl: 116.5±4.73%, Hyase: 100.4±5.3%, p=0.01; Ch-ABC, 112.0±4.0%; p=0.5; Figs. 5B). Overall, our results show a crucial role of hyaluronic acid in the molecular mechanisms of iLTP induction in PV→PC synapses, but did not support a significant contribution of CSPGs to the plastic changes in these synapses.

**Figure 5.**
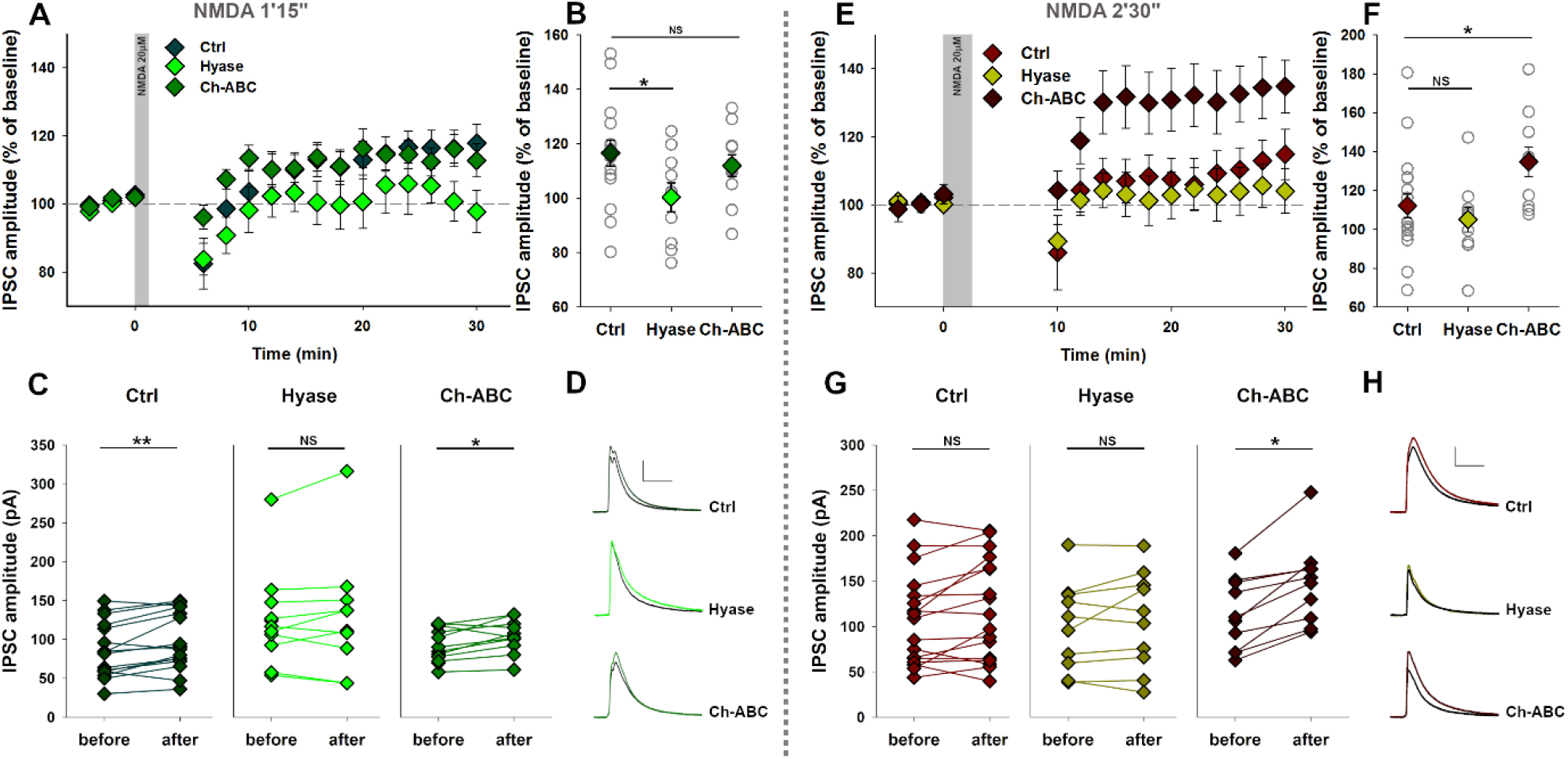
Chondroitin sulfate and hyaluronic acid degradation affect NMDA-induced GABAergic plasticity in inhibitory synapses between CA1 parvalbumin (PV) interneurons and pyramidal cells (PC) **A**. GABAergic long-term potentiation (iLTP) induced at PV→PC synapses by bath application of NMDA for 1 minute 15 seconds, recorded in sham-treated slices (Ctrl), and slices treated for 2 hours with hyaluronidase (Hyase) or chondroitinase ABC (Ch-ABC). The gray area indicates the duration of NMDA application. **B**. Summary plot of iLTP magnitudes assessed 24-30 minutes after plasticity induction in PV→PC synapses by NMDA application for 1 minute 15 seconds (Ctrl, n=16; Hyase, n=11; Ch-ABC, n=11). Diamonds represent mean ± SEM. Statistical significance was determined by t-test vs. Ctrl group. **C**. Comparison of IPSC amplitudes in the PV→PC synapses recorded before and 26-30 minutes after iLTP induction with NMDA (applied for 1 min 15 s). Statistical significance was determined using the paired t-test. **D**. Representative traces showing IPSCs at the PV→CA1 PC synapses recorded before (black) and 25-30 minutes after iLTP induction (green) in control (top), hyaluronidase-treated (middle), and chondroitinase-treated (bottom) slices (scale: 40 pA, 40 ms). **E**. GABAergic long-term plasticity induced at PV→PC synapses by bath application of NMDA for 2 minutes 30 seconds, recorded in sham-treated slices (Ctrl), and slices treated for 2 hours with hyaluronidase (Hyase) or chondroitinase ABC (Ch-ABC). The gray area indicates the duration of NMDA application. Note the lack of PV→PC plasticity in the sham-treated controls. **F**. Summary plot of iLTP magnitudes assessed 24-30 minutes after plasticity induction in PV→PC synapses by NMDA application for 2 minutes 30 seconds (Ctrl, n=18; Hyase, n=10; Ch-ABC, n=10). Diamonds represent mean ± SEM. Statistical significance was determined by t-test vs. Ctrl group. **G**. Comparison of IPSC amplitudes in PV→PC synapses recorded before and 26-30 minutes after NMDA application for 2 minutes 30 seconds. Statistical significance was determined using the paired t-test. **H**. Representative traces showing IPSCs at the PV→CA1 PC synapses recorded before (black) and 25-30 minutes after NMDA application for 2 min 30 s (brown/green) in control (top), hyaluronidase-treated (middle), and chondroitinase-treated (bottom) slices (scale: 40 pA, 40 ms). *p < 0.05, **p < 0.01, ns – not significant

Given that enzymatic cleavage of hyaluronic acid in brain slices reduces dendritic Ca^2+^ influx [5], we investigated whether extending the duration of NMDA application from 1 min 15 s to 2 min 30 s could rescue the impaired iLTP observed in HA-depleted slices by prolonging membrane depolarization. Surprisingly, NMDA application for 2 min 30 s does not induce significant long-term plasticity in the CA1 PV→PC synapses in sham-treated control slices (IPSC amplitude: 106.6±11.9 pA before, 112.7±12.2 pA after 2 min 30 s NMDA, p=0.33, paired *t*-test; Figs. 5G-I). Similarly, we observed a lack of GABAergic plasticity after prolonged NMDA exposure in slices treated with hyaluronidase (100.3±15.4pA before, 106.5±14.2pA after, p=0.29; Figs. 5G-I). Conversely, in slices where chondroitin sulfate was digested, NMDA application for 2 min 30 s induced significant stable IPSC potentiation (113.1±12.6pA before, 147.6±14.2pA after, p=0.0007, paired *t*-test; Fig. 5G). This indicates that both the sham-treated control slices and those treated with hyaluronidase showed no iLTP in response to prolonged NMDA application (change in IPSC amplitude, Ctrl: 108.2±6.5%, Hyase: 105.0±6.4%, p=0.46). In contrast, digestion of CSPGs elicited a significant iLTP in PV→PC synapses using the same protocol (Ch-ABC: 134.8±7.7%, p=0.03 vs. Ctrl; Figs. 5E,F). Furthermore, the magnitude of iLTP induced by the prolonged NMDA application in Ch-ABC-treated slices was significantly greater than that induced by shorter NMDA application in control conditions (Fig. S3). Thus, the recovery of iLTP by the adjustment of the NMDA administration protocol confirms the involvement of chondroitin sulfate-containing ECM elements in the modulation of PV→PC synaptic strength. These findings also suggest that local modification of brain CSPGs can unveil a latent capacity for GABAergic plasticity induction.

## Discussion

In this study, we demonstrate that enzymatic digestion of chondroitin sulfate proteoglycans (CSPGs) and hyaluronic acid (HA) exerts differential effects on GABAergic synaptic plasticity in the hippocampal CA1 region. Specifically, CSPG degradation impairs NMDA-induced inhibitory long-term potentiation (NMDA-iLTP) at dendrite-targeting SST→CA1 PC synapses, while leaving PV→PC synapses unaffected (Fig. 3). Conversely, hyaluronic acid is crucial for iLTP induction at PV→PC synapses (Fig. 5). Intriguingly, while prolonged NMDA application (2 min 30 sec) did not induce iLTP at PV→PC synapses under control conditions, CSPG degradation enabled iLTP induction after long NMDA treatment, suggesting that a form of cryptic inhibitory plasticity in this system may depend on the ECM state. These findings underscore the synapse-specific contributions of the ECM to synaptic GABAergic plasticity.

### Impact on basal inhibitory synaptic transmission

Despite the effects of ECM digestion on NMDA-induced inhibitory plasticity, no changes were observed in basal IPSCs from either SST-or PV-positive interneurons upon CSPG or HA degradation (Fig. 1). This indicates that ECM degradation specifically influences GABAergic plasticity without altering the basal strength of inhibitory synapses. However, some studies have reported that Ch-ABC treatment can downregulate inhibitory synaptic transmission, particularly from PV interneurons, or reduce the number of GABAergic synapses on PV interneurons [10], [12], [31]. If ECM degradation affects basal inhibitory synaptic transmission, it may involve other inhibitory inputs to the CA1 pyramidal cells, distinct from SST+ or PV+ interneurons. These inputs could potentially originate from cholecystokinin-positive (CCK) interneurons, whose synaptic transmission is modulated by dystroglycan, a membrane receptor interacting with the ECM [32], [33]. Additionally, PV- and SST-positive interneurons encompass various subgroups, such as bistratified cells, O-LM cells, PV basket cells, and axo-axonic interneurons [34], which may be differentially affected by ECM modification. Furthermore, the absence of significant changes in the rise and decay kinetics of IPSCs after ECM degradation suggests that such treatment does not alter the synaptic subunit composition of GABA_A_ receptors [35] or the extracellular environment that shapes synaptic GABA transients [36], [37], [38], [39].

### GABAergic short-term plasticity and ECM integrity

ECM integrity is known to influence the lateral diffusion of glutamate receptors within both the synaptic and extrasynaptic regions of the neuronal membrane. Previous studies have shown that CSPG digestion increases the lateral mobility of AMPA receptors, leading to faster exchange of desensitized synaptic receptors with naïve receptors from extrasynaptic membranes [27]. Consequently, Ch-ABC treatment affects short-term plasticity at excitatory synapses, as evidenced by changes in synaptic responses to paired-pulse stimulation. Similarly, application of the extracellular matrix metalloprotease MMP9 affects the membrane mobility of synaptic NMDA receptors [40]. However, in our study we did not observe changes in the short-term plasticity of GABAergic transmission, assessed by paired-pulse stimulation, following ECM degradation (Fig. S1). This contrasts with recent studies indicating that the application of MMP3, another extracellular metalloproteinase [41], [42], induces iLTP and slows the membrane mobility of GABA receptors containing the α1 subunit [29]. Notably, this effect was observed only in the synaptic, not extrasynaptic, regions of GABA receptor mobility trajectories.

Given that synaptic transmission leads to significant desensitization of GABA_A_ receptors [43] and their enhanced diffusion out of inhibitory synapses [44], it is clear that the exchange of synaptic GABA_A_ receptors is tightly regulated [45]. Therefore, to further investigate whether ECM degradation might cause changes in short-term plasticity, we applied prolonged burst stimulation to inhibitory synapses. This approach revealed deepened burst-induced short-term synaptic depression in slices treated with either Hyase or Ch-ABC. This effect was significant only at PV→PC synapses, suggesting that short-term plasticity at distinct types of inhibitory synapses is differentially affected by ECM degradation. Additionally, PV→PC synapses also exhibited more pronounced paired-pulse depression and burst-induced depression than dendrite-targeting SST input, suggesting different subunit compositions of GABA_A_ receptors in these synapses (Fig. 2).

### ECM degradation impairs inhibitory plasticity

The impact of PNN modification or degradation on long-term synaptic plasticity at excitatory synapses is diverse and varies significantly depending on the brain region and type of PNN manipulation involved [4]. In the hippocampal CA1 region, several studies have shown that PNN digestion leads to a reduction in LTP [5], [24], [46], [47]. However, the effects of ECM degradation are not uniform and can vary depending on the synapse type and the brain region studied. For example, in the CA2 region, which is crucial for social memory and typically does not exhibit LTP, ECM digestion facilitates excitatory LTP induction [48]. Similarly, ECM degradation with Ch-ABC enhances excitatory long-term depression (LTD) [49], [50]. These findings underscore the critical role of PNN integrity and the specific molecular composition of the ECM in maintaining excitatory synaptic plasticity in a synapse-specific manner.

Long-term inhibitory plasticity often involves heterosynaptic induction mechanisms, meaning it is induced by plastic changes in neighbouring excitatory synapses [51], [52]. Consequently, plastic changes in GABAergic synapses are often coexpressed with excitatory plasticity. Thus, our finding that ECM degradation affects inhibitory plasticity induced by brief NMDA application, which is also known to induce excitatory plasticity, is not surprising (Figs. 3, 5). However, studies on inhibitory plasticity must consider the diversity of interneuron types, each with distinct network roles and synaptic properties [34]. Our study highlights synapse-specific roles of the brain ECM in GABAergic plasticity. CSPG degradation impaired iLTP at synapses formed by dendrite-targeting SST-positive interneurons, whereas iLTP at PV→PC synapses was compromised by hyaluronan digestion. This suggests that different ECM components are critical for different types of inhibitory synaptic plasticity. Notably, HA is involved in GABAergic plasticity at synapses located in the soma and proximal dendrites, where HA-rich PNNs are prominent [5]. Additionally, despite the partially overlapping substrate specificity of hyaluronidase and chondroitinase [53], our results indicate divergent effects of ECM digestion, underscoring synapse-specific roles of HA and CSPGs in GABAergic plasticity.

### Potential role of integrins

The brain ECM is a source of bioactive compounds released after extracellular proteolysis or enzymatic cleavage of glycosaminoglycans, which can affect adhesion proteins [54] or membrane calcium channels [55]. Integrins, which act as major receptors for ECM components, are crucial regulators of synaptic transmission [56]. For instance, activation of β1 integrins enhances dendritic spine motility following CSPG digestion [22]. Additionally, β1 or β3 subunit-containing integrins control the dynamics of glycine receptors and the scaffolding protein gephyrin at the glycinergic synapses [28]. Recently, our group has revealed that integrins modulate GABAergic synaptic transmission and plasticity. Specifically, interference with integrin-dependent adhesion blocks the induction of NMDA-iLTP in CA1 pyramidal cells or interneurons [20], [57]. However, we did not observe changes in levels of extended, presumably active, forms of integrin β1 or β3 (Fig. 4). This lack of changes in active integrin levels after Hyase or Ch-ABC treatment suggests that integrin-dependent adhesion and signaling are not involved in the impairment of NMDA-iLTP following ECM degradation.

### CSPGs digestion unveils cryptic GABAergic plasticity

Our study reveals that the digestion of chondroitin sulfate proteoglycans (CSPGs) unveils cryptic GABAergic plasticity at PV→PC synapses induced by prolonged NMDA application for 2 min 30 sec (Fig. 5). This observation parallels the enhanced magnitude or facilitation of excitatory LTD observed in slices treated with Ch-ABC [49], [50]. Such opposing effects on inhibitory and excitatory plasticity can significantly alter the input-output characteristics of local neuronal circuits. In this context, it is worth mentioning that inhibitory LTD at PV→PC synapses in the hippocampal CA2 region emerges with the maturation of perineuronal nets and is impaired following CSPG degradation by Ch-ABC [58]. These findings highlight the critical role of CSPGs in constraining both excitatory and inhibitory synaptic plasticity, and suggest that cryptic iLTP at PV→PC synapses could play a significant role in modulating neural circuit dynamics and memory processes.

### Conclusions and future directions

Inhibitory plasticity is primarily thought to regulate memory precision [59]. Our findings underscore the crucial role of CSPGs in constraining inhibitory synaptic plasticity at PV→PC synapses, which may be essential for maintaining this precision. Ramsaran et al. [17] reported that in juvenile mice, an immature ECM in the hippocampus leads to imprecise memory due to weaker synaptic transmission from PV interneurons and incorporation of more neurons into the engram. Notably, the functional maturation of PV-expressing interneurons in CA1, facilitated by the assembly of mature ECM, is necessary for sparse engram formation and memory precision [17]. Taking this into account, it may be expected that future research should explore how local ECM modifications, by altering the GABAergic plasticitome [60], can affect memory formation in adult animals.

Interference with ECM integrity or its endogenous cleavage may result in memory enhancement. For instance, digestion of PNNs is associated with improved recognition memory [49], [50]. Similarly, knockout mice lacking tenascin-R exhibit significantly improved working memory and reversal learning in the Morris water maze task [61]. Additionally, brain-wide knockout of aggrecan, which leads to PNN ablation, improves recognition memory in mice [62]. Moreover, MMP3 knockout mice demonstrate impaired iLTP, enhanced contextual fear conditioning, and faster learning in the Morris water maze [29]. Given that PNN manipulation can reopen the critical period of enhanced plasticity and learning capacity [63], [64], future studies should investigate how synapse-specific interactions between inhibitory plasticity and ECM can be engaged in this process. Understanding the interplay among all the constituents of plastic inhibitory tetrapartite synapses could provide new insights into the molecular and network mechanisms of learning and memory.

## Acknowledgments

This work was supported by funding from the National Science Centre (Poland) SONATA 2017/26/D/NZ4/00450 (to G.W.); J.J and J.W.M. were supported by National Science Centre (Poland) grant OPUS 2021/43/B/NZ4/01675.

