## Supplement for "Extracellular matrix integrity regulates GABAergic plasticity in the hippocampus"

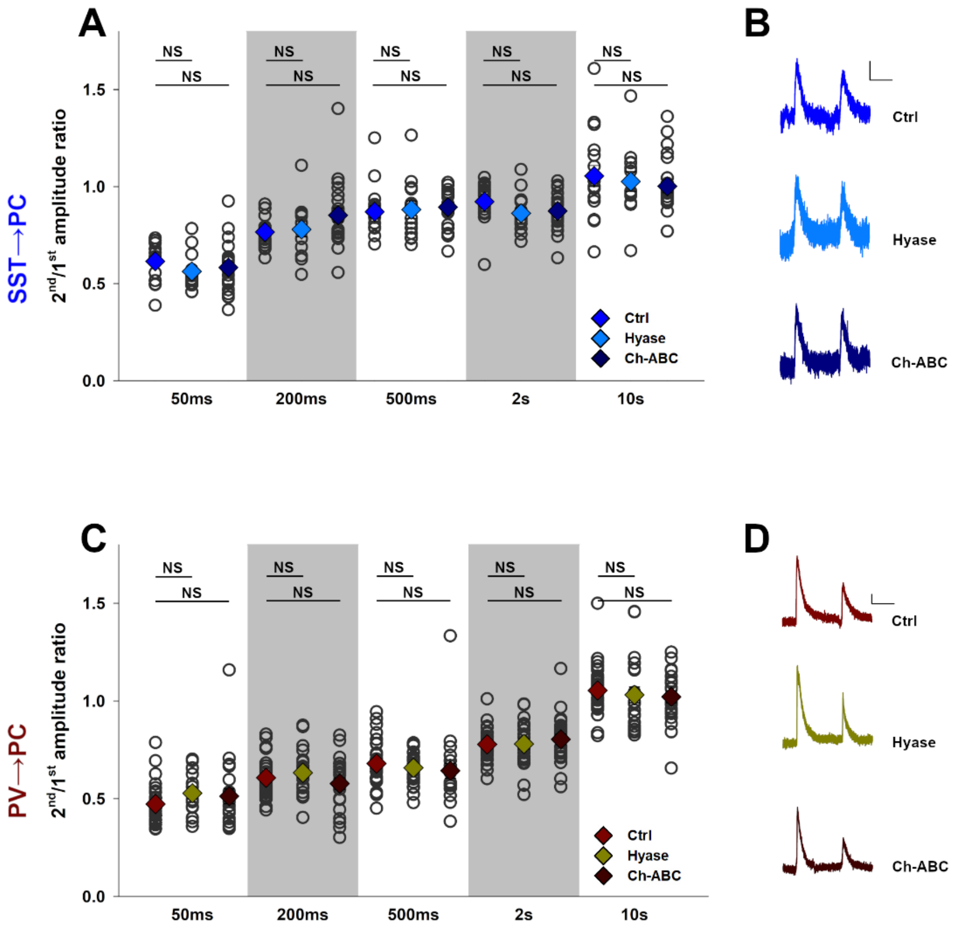


***Supplemental Figure 1. Effects of enzymatic degradation of hyaluronic acid and chondroitin sulfate proteoglycans on short-term plasticity of inhibitory synaptic transmission in the CA1 field of the hippocampus***

***A.*** *Short-term plasticity in SST→CA1 pyramidal cell (PC) synapses was assessed using paired-pulse stimulation applied at different intervals. Paired-pulse ratios (amplitude of the 2nd IPSC divided by the amplitude of the 1st response in pairs) were recorded in the SST→PC synapses at stimulation intervals of 50 ms, 200 ms, 500 ms, 2 s, and 10 s. Diamonds represent mean ± SEM. Statistical significance was determined by t-test vs. Ctrl group (50ms: Ctrl: 0.62±0.02, Hyase: 0.56±0.03 p=0.16, Ch-ABC: 0.58±0.03 p=0.41; 200ms: Ctrl: 0.77±0.02, Hyase: 0.78±0.04 p=0.74, Ch-ABC: 0.85±0.04 p=0.06; 500ms: Ctrl: 0.87±0.03, Hyase: 0.89±0.04 p=0.84, Ch-ABC: 0.89±0.02 p=0.52; 2s: Ctrl: 0.92±0.03, Hyase: 0.86±0.03 p=0.12, Ch-ABC: 0.87±0.02 p=0.15; 10s: Ctrl: 1.05±0.05, Hyase: 1.03±0.05 p=0.71, CHh-ABC: 1±0.03 p=0.39). ns – not significant*

***B.*** *Representative traces of paired-pulse responses to stimulation with a 200 ms interval recorded in SST→PC synapses in Ctrl (top), Hyase (middle), and Ch-ABC (bottom) treated slices (scale: 10 pA, 100 ms).*

***C.*** *Short-term plasticity in the PV→CA1 pyramidal cell (PC) synapses was assessed using paired-pulse stimulation applied at different intervals. The paired-pulse ratios are shown. Diamonds represent mean ± SEM. Statistical significance was determined by t-test vs. Ctrl group: (50ms: Ctrl: 0.47±0.02, Hyase: 0.53±0.02 p=0.05, Ch-ABC: 0.51±0.04 p=0.28; 200ms: Ctrl: 0.61±0.02, Hyase: 0.63±0.03 p=0.4, Ch-ABC: 0.58±0.03 p=0.36; 500ms: Ctrl: 0.68±0.02, Hyase: 0.66±0.02 p=0.45, Ch-ABC: 0.64±0.04 p=0.35; 2s: Ctrl: 0.78±0.01, Hyase: 0.78±0.03 p=0.93, Ch-ABC: 0.8±0.03 p=0.36; 10s: Ctrl: 1.05±0.02, Hyase: 1.03±0.04 p=0.6, Ch-ABC: 1.02±0.03 p=0.37). ns – not significant*

***D.*** *Representative traces of paired-pulse responses to stimulation with a 200 ms interval recorded in PV→PC synapses in Ctrl (top), Hyase (middle), and Ch-ABC (bottom) treated slices (scale: 10 pA, 100 ms).*

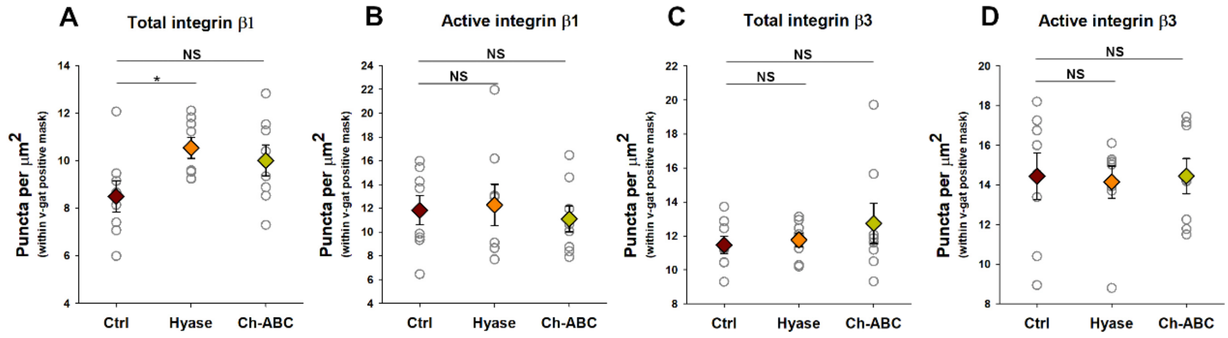


***Supplemental Figure 2. Impact of hyaluronan and chondroitin sulphate degradation on the density of total pool and active forms of integrin β1 and β3 in CA1 stratum radiatum.***

***A-D**. Analysis of the density of integrin puncta near inhibitory synapses identified by vGAT staining, in the CA1 stratum radiatum. The figures show data for: total pool of integrin β1 (A)**,** active integrin β1 (B), total pool of integrin β3 (C), and active integrin β3 (D).*

*Data represent n=8 independently treated slices from 4 mice. Colored diamonds indicate mean ± SEM. Statistical significance was determined by t-test vs. Ctrl group: *p < 0.05, ns – not significant.*

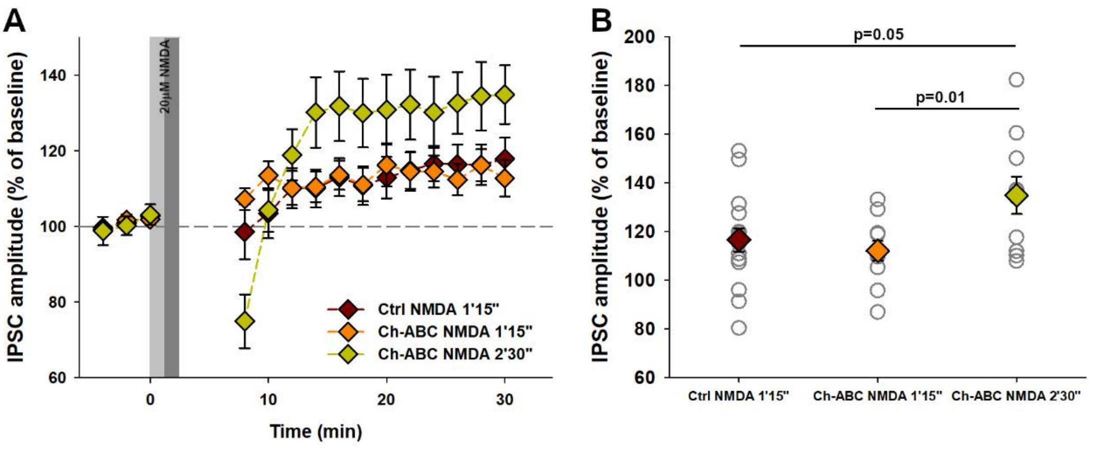


***Supplemental Figure 3. Comparison of iLTP induced by different protocols in sham-treated and chondroitinase-treated slices.*** *The figure contains the data already presented in Figs. 5A, B, E, and F.*

***A**. GABAergic iLTP induced at PV→PC synapses by bath application of NMDA for 1 minute 15 seconds (1'15") in sham-treated slices (brown) and chondroitinase ABC-treated slices (orange), as well as NMDA application for 2 minutes 30 seconds (2'30") in chondroitinase ABC-treated slices (green). The gray areas indicate the two different durations of NMDA application.*

***B.** Summary plot of iLTP magnitudes assessed 24-30 minutes after plasticity induction in PV→PC synapses by NMDA application for 1 minute 15 seconds (Ctrl NMDA 1'15": 116.5 ± 4.7%, n=16; Ch-ABC NMDA 1'15": 112.0 ± 4.0%, n=11; Ch-ABC NMDA 2'30": 134.8 ± 7.7%, n=10). Diamonds represent mean ± SEM. Statistical significance was determined by the t-test, with p-values presented above the horizontal bars.*
